# The Omicron (B.1.1.529) SARS-CoV-2 variant of concern also affects companion animals

**DOI:** 10.1101/2022.04.13.488132

**Authors:** Lidia Sánchez-Morales, José M. Sánchez-Vizcaíno, Marta Pérez-Sancho, Lucas Domínguez, Sandra Barroso-Arévalo

## Abstract

The recent emergence of the Omicron variant (B.1.1.529) has brought with it a large increase in the incidence of SARS-CoV-2 disease worldwide. However, there is hardly any data on the incidence of this new variant in companion animals. In this study, we have detected the presence of this new variant in domestic animals such as dogs and cats living with owners with COVID19 in Spain that have been sampled at the most optimal time for the detection of the disease. None of the RT-qPCR positive animals (10.13%) presented any clinical signs and the viral loads detected were very low. In addition, the shedding of viral RNA lasted a short period of time in the positive animals. Infection with the Omicron variant of concern (VOC) was confirmed by a specific RT-qPCR for the detection of this variant and by sequencing. These outcomes suggest a lower virulence of this variant in infected cats and dogs. This study demonstrates the transmission of this new variant from infected humans to domestic animals and highlights the importance of doing active surveillance as well as genomic research to detect the presence of VOCs or mutations associated with animal hosts.

## Introduction

The pandemic associated with the COronaVIrus Disease 2019 (COVID-19), produced by the SARS-CoV-2 virus, has been active for almost two years now. From the beginning, 406 million cases have been confirmed in the world, with 6.09 million deaths according to the last World Health Organization (WHO) report (World Health Organisation, 2022).

The SARS-CoV-2 is an RNA virus whose organization is shared with other *Beta* coronaviruses. The genome of this virus consists of 13 opening reading frames (ORFs) and 15 non-structural proteins (NSP). The ORFs, from 5’ to 3’, codify for replicase (ORF1a/ORF1b), spike (S) protein, envelope (E) protein, membrane (M) protein, and nucleocapsid (N) protein (Hu, Guo et al. 2021, Raj 2020; Viralzone, 2022). Due to these genomic characteristics, this coronavirus has been suffering a great number of mutations, mainly in the spike protein, which may have influenced the virus transmission rate, the disease severity, or abrogate the immunity produced by the vaccines, among other factors (Zou, Xia et al. 2021). The apparition of these mutations has triggered virus evolution. Consequently, new variants have been identified. According to the pathogenic potential and the virulence of the different isolates, the WHO have classified them into variants of concern (VOCs), variants of interest (VOIs), and variants under monitoring (VUMs) (He, Hong et al. 2021). Until December 2021, four VOCs had been reported: Alpha (B.1.1.7), Beta (B.1.351), Gamma (P.1), and Delta (B.1.617.2) (Saxena, Kumar et al. 2021). On November 26^th^, a new variant was determined by the WHO as the 5^th^ VOC, named Omicron (B.1.1.529). The first sample identified as this variant was taken in South Africa on the 9^th^ of November, while the first sequenced case was from a sample collected in Botswana on the 11^th^ of November (He, Hong et al. 2021).

Until now, the Omicron variant is the VOC with the largest number of mutations detected, with 34 of them accumulated in the spike protein. Several of these mutations in the spike protein have been related to increased viral antibody neutralization evasion capacity or higher affinity between the spike/ACE receptor binding (Zou, Xia et al. 2021), facilitating the virus entry into the cell. Thus, this constellation of mutations appears to have influenced virus transmissibility, severity, and immune evasion (CDC 2021, He, Hong et al. 2021, Araf, Akter et al. 2022). These mutations have led to greater contagiousness than the previous variants as well as different clinical signs, which consist of slight fever, myalgia, fatigue, and shortness of breath. However, the most dangerous characteristic of this variant is its high rate of immune escape even in previously immunized by natural infection and vaccinated people (Meo, Meo et al. 2021). Because of these characteristics, the Omicron variant has gained great concern in public health worldwide.

In Spain, according to data published by the Ministry of Health, the cumulative incidence of SARS-CoV-2 rose from 77 cases per 100,000 inhabitants on 15^th^ November 2021 to 465 on 15^th^ December 2021, showing a significant increase. It continued growing until reaching 3,418 on January 20^th^, 2022. This growth in cases coincided with the dates on which the new Omicron variant was first detected in Spain, around mid-December 2021. As could be expected, sequence analysis since that period has revealed the growing dominance of the new VOC in the country (Ministerio de Sanidad de España, 2021).

Within the Omicron variant, three lineages or subvariants are distinguished so far: B.1.1.529.1 (BA.1), B.1.1.529.2 (BA.2), and B.1.1.529.2 (BA.3). BA.2 lineage has 32 mutations shared with BA.1, but 28 distinct, and BA.3 spike protein is a combination of BA.1 and BA.2 with no new mutations. The most prevalent is the BA.1 sublineage, while BA.2 has been observed to reinfect patients previously infected with BA.1, being more prevalent in Denmark (Chen and Wei 2022; Mohapatra, Kandi et al. 2022; Desingu and Nagarajan, 2022). Shortly after the SARS-CoV-2 virus entered our lives, field studies on the incidence in animals, as well as experimental studies, began to be carried out to learn about their role in this new disease (Shi, Wen et al. 2020). In the case of animals, several studies have found that some species such as *Canis lupus familiaris, Nevison vison, Manis javanica, Mesocricetus auratus* and *Odocoileus virginianus* are susceptible to the Omicron variant, as revealed by sequencing results from natural or experimental infection (GISAID). However, experimental studies on hamsters have evidenced the lower pathogenicity of this variant in comparison with Delta and B.1.1 variants in this species, based on different variables such as body weight and respiratory function (Suzuki, Yamasoba et al. 2022). The conclusions obtained by these studies were that, unlike other VOCs, Omicron is not able to efficiently replicate in the lower respiratory tract of Syrian hamsters, which results in the detection of lower viral loads and fewer pathology findings in the lungs of the experimentally infected animals comparing with infection by other isolates (Abdelnabi, Foo et al. 2022). In addition, inoculated mice with B.1.1.529 (Omicron) had lower levels of pro-inflammatory cytokines and chemokines, on occasions similar to non-infected mice, than those inoculated with the B.1.351 (Beta) variant (Halfmann et al. 2022).

Despite these results suggesting a lower virulence of this variant in infected animals, very different results were observed in an experimental study in wild carnivores (mink, *Neovison vison*) which are known to be very susceptible to SARS-CoV-2 virus infection. In this study, minks were infected with the Omicron variant. The animals became ill, had clinical symptoms, positive PCR results, as well as macroscopic and microscopic lesions post mortem (Virtanen, Aaltonen et al. 2022). All these aspects lead us to wonder what will be the relevance of the infection with the Omicron variant in species in close contact with humans such as dogs and cats in comparison with the previously described VOCs (Barroso-Arevalo, Rivera et al. 2021, Barroso-Arévalo, Sánchez-Morales et al. 2022, Doerksen, Lu et al. 2021). Is transmissibility to susceptible pets higher with this variant, as is occurring in the case of humans? What are the clinical repercussions of the infection in cats and dogs? To elucidate the implications of infection with the Omicron variant in pets, we have carried out an active sampling of cats and dogs in close contact with SARS-CoV-2 infected people with clinical signs compatible with this variant and/or confirmed by RT-qPCR or sequencing. In this study, we have observed a low prevalence of infection in the animals, as well as low viral loads in the positive cases, despite the samplings being carried out at the optimum moment to detect human-to-pet transmission.

## Material and methods

### Animal and owner sample collection

Samples from domestic animals including cats (n=28), dogs (n=50), and rabbit (n=1) were taken between the 15^th^ of December 2021 to 24^th^ of March 2022. A total of 69 animals (21 cats and 47 dogs) were from Madrid, 6 animals (3 cats and 3 dogs) from Galicia, and 4 cats (from the Basque Country). All these animals were sampled during the quarantine period of their owner and, therefore, had been in contact with positive people for SARS-CoV-2. The samples were taken using protocols approved by the Complutense University of Madrid’s Ethics Committee for Animal Experiments (Project License 14/2020). Owners were informed about the purpose of the study as well as the data protection policy. When possible, samples were taken on consecutive days to gather more information about the potential animal infection. The samples consisted of oral/nasal and rectal swabs collected in DeltaSwab® Virus containing 3ml of viral transport media (MTV) (Deltalab S.L., Cataluña, Spain) and sera if possible that were collected in tubes without anticoagulant. All the samples were refrigerated and taken to the Health Surveillance Centre (VISAVET) at the Complutense University of Madrid and stored at -80ºC until analysis. In addition, a survey of the owners was carried out in order to know the potential symptoms they were presenting to confirm Omicron variant associated signs, as well as a nasal swab sample collection in some cases to confirm the SARS-CoV-2 variant involved in the infection by RT-qPCR and sequencing.

### Detection of SARS-CoV-2 infection by reverse transcription-quantitative PCR (RT-qPCR) and specific Omicron RT qPCR and virus isolation

Total RNA was extracted using the column-based High Pure Viral Nucleic Acid Kit (Roche, Basel, Switzerland), according to the manufacturer’s instructions. Total RNA was suspended in RNase/DNase-free water and stored at -80ºC. The detection of the RNA of SARS-CoV-2 was carried out using a diagnostic RT-qPCR, hereafter “Diagnosis PCR”, based on the detection of the envelope protein (E)-encoding gene (Sarbeco) and two targets (IP2 and IP4) of the RNA-dependent RNA polymerase gene (RdRp) in an RT-qPCR protocol established by the World Health Organization according to the guidelines that can be found at https://www.who.int/emergencies/diseases/novel-coronavirus-2019/technical-guidance/laboratory-guidance (Corman, Landt et al. 2020).

An specific RT-qPCR was used for the identification of the SARS-CoV-2 Omicron variant, hereafter “Omicron PCR”, targeting both the envelope protein (E) - encoding gene as well as an Omicron-specific spike insertion-deletion mutation (indel_211-214) found in the B.1.1.529/BA.1 lineage and BA.1.1 sublineage, so in the case of the BA.2 and BA.3 Omicron lineages would only be detected by the gen E target. The kit used was the SuperScript III Platinum One-Step qRT-PCR kit (Invitrogen) according to the protocol described in (Sibai, Wang et al. 2022).

Positive samples for RT-qPCR were subjected to attempts of viral isolation using the previously described methods in (Gortázar, Barroso-Arévalo et al. 2021).

### Whole-genome sequencing and phylogenetic analysis

Whole-genome sequences were obtained from the two positive oropharyngeal swabs samples with the higher viral loads based on Ct values (Ct of 32.45 and 30.01) by both “Diagnosis” and “Omicron” RT-qPCRs, following the protocol described by (Paden, 2020). Sequence analysis was performed using the Sequencing Analysis software v.5.3.1(Applied Biosystems), while SeqScape v.2.5 software (Applied Biosystems) was used for sequence assembly using the SARS-CoV-2 isolate Wuhan-Hu-1, complete genome (GenBank accession number: NC_045512) as a reference genome.

Phylogenetic analysis was performed using MEGA X software (Tamura, 1992). Four sequences were obtained from this study (Dog_ 8, Cat_19, Owner_1, and Owner_2, which correspond with one dog, one cat, the dog’s owner, and the owner of Cat_26, 27, and 28. Unfortunately, no positive sample for sequencing was available from the owner of Cat_19. In the case of cats 26, 27, and 28, sequencing was not possible because of the low RNA loads of the positive samples (Table 1).

**Table 1.** Positive animals to RT-qPCR including specie, date of sampling, sample type, the day post-infection of the owner (DPI), RT-qPCR Cycle threshold (Ct) value in both “Diagnosis PCR” and “Omicron PCR”, sequence if available, and viral isolation result.

| Animal, date of collection | Sample type | DPI owner | Diagnosis RT-qPCR Ct | Omicron RT-qPCR Ct | Sequence | Viral isolation |
| --- | --- | --- | --- | --- | --- | --- |
| Cat_2, 20th December, 2021 | Rectal swab | 2 DPI | 34.2 | 36.85 | NA | Negative |
| Dog_8, 1<sup>st</sup> January, 2022 | Oropharyngeal swab | 3 DPI | 32.45 | 36.92 | B.1.1.529 | Negative |
| Cat_7, 16<sup>th</sup> January, 2022 | Oropharyngeal swab | 2 DPI | 36.6 | 38.03 | NA | Negative |
| Cat_19, 22<sup>th</sup> January, 2022 | Oropharyngeal swab | 3 DPI | 30.01 | 32.78 | B.1.1.529 | Negative |
| Cat_13, 27<sup>th</sup> January, 2022 | Oropharyngeal swab | 4 DPI | 35.7 | 37.67 | NA | Negative |
| Cat_26, 18<sup>th</sup> March, 2022 | Oropharyngeal swab | 2 DPI | 34.5 | 39.65 | NA | Negative |
| Cat_27, 18<sup>th</sup> March, 2022 | Oropharyngeal swab | 3 DPI | 35.6 | 39.05 | NA | Negative |
| Cat_28, 19<sup>th</sup> March, 2022 | Oropharyngeal swab | 4 DPI | 35 | 37.86 | NA | Negative |

\*NA: Not available

A total of 31 additional representative sequences were used for the analysis, including sequences from cats and dogs, the reference genome from Wuhan, as well as variants of concern such as the B.1.1.7 variant from the United Kingdom, variant B.1.35 from South Africa, variant B.1.617.2 from India, variant B.1.1.248 from Brazil and lineages BA.1 and BA.2 of the B.1.1.529 Omicron variant.

The final alignment involved 35 whole-genome sequences with an average amino acid p-distance (1-amino acid identity) lower than 0.001, which is considered adequate since it is within the acceptance threshold of <0.8 (Tamura, 1992). This alignment was used to build the phylogenetic tree using the maximum likelihood method and bootstrap testing of 2,000 replicates. The best model was the Tamura-Nei Model, so it was the one used to create the phylogenetic tree.

### Virus neutralization test (VNT) for detection of specific neutralizing antibodies against SARS-CoV-2

Virus neutralization test (VNT) was used to confirm the presence of neutralizing antibodies against SARS-CoV-2 in all the sera collected.

Briefly, the VNT was performed in duplicate in 96-well plates by incubating 25 μL of two-fold serially diluted sera with 25 μL of 100 TCID_50_/ml of SARS-CoV-2. The virus-serum mixture was incubated at 37°C with 5% CO_2_. At 1-hour post-incubation, 200 μL of Vero E6 cell suspension were added to the virus-serum mixtures, and the plates were incubated at 37°C with 5% CO_2_. The neutralization titers were determined at 3 days post-infection. The titer of a sample was recorded as the reciprocal of the highest serum dilution that provided at least 100% neutralization of the reference virus, as determined by the visualization of cytopathic effect (CPE). In addition, at the end of the period (3 days post-infection), cells were fixed with 6% paraformaldehyde and then stained with crystal violet to observe the cytopathic effect.

## Results

### Sampling data

A total of 50 dogs, 28 cats, and 1 rabbit (n=79) were sampled during the quarantine period of their owners from 15th December 2021 to 23^h^ March 2022, coinciding with the period of the highest prevalence of the Omicron variant in Spain (Ministerio de Sanidad de España., 2022) in Madrid (n=69), Galicia (n=6) and the Basque Country (n=4).

All the owners reported a high level of contact with the pets included in the sampling during their confinement period. None of the animals showed any compatible symptoms with the illness.

### SARS-CoV-2 infection prevalence

SARS-CoV-2 RNA was detected by RT-qPCR in seven cats and one dog by both “Diagnosis PCR” and “Omicron PCR”. This represents 10.13% of the total analyzed animals. All of the positive animals were sampled in Madrid. All the positive samples for RT-qPCR were negative for viral isolation (Table 1).

### Neutralizing antibodies detection by VNT

Sera were collected from 15 animals (1 cat and 14 dogs), including Dog_8 and Cat_13 which were also positive for RT-qPCR (15 and 20 days after RT-qPCR positive result, respectively). Among the 15 serum samples, none of them presented neutralizing antibodies.

### Whole-genome sequencing and phylogenetic analysis

The complete genome sequence of SARS-CoV-2 was obtained from the oropharyngeal swabs from both Dog_8 and Cat_19 (GenBank accession numbers: ON115270 and ON115269; GISAID accession ID: EPI_ISL_11580532 and EPI_ISL_11580576) as well as from nasopharyngeal swabs from the owner of Dog_8 (Owner_1; GenBank accession numbers: ON115271; GISAID accession ID: EPI_ISL_11580604) and the owner of Cat_26, 27 and 28 (Owner_2: GenBank accession number: ON115272; GISAID accession ID: EPI_ISL_11580636) since the remaining PCR-positive animals had too low viral RNA loads for effective sequencing.

**Figure 1.**
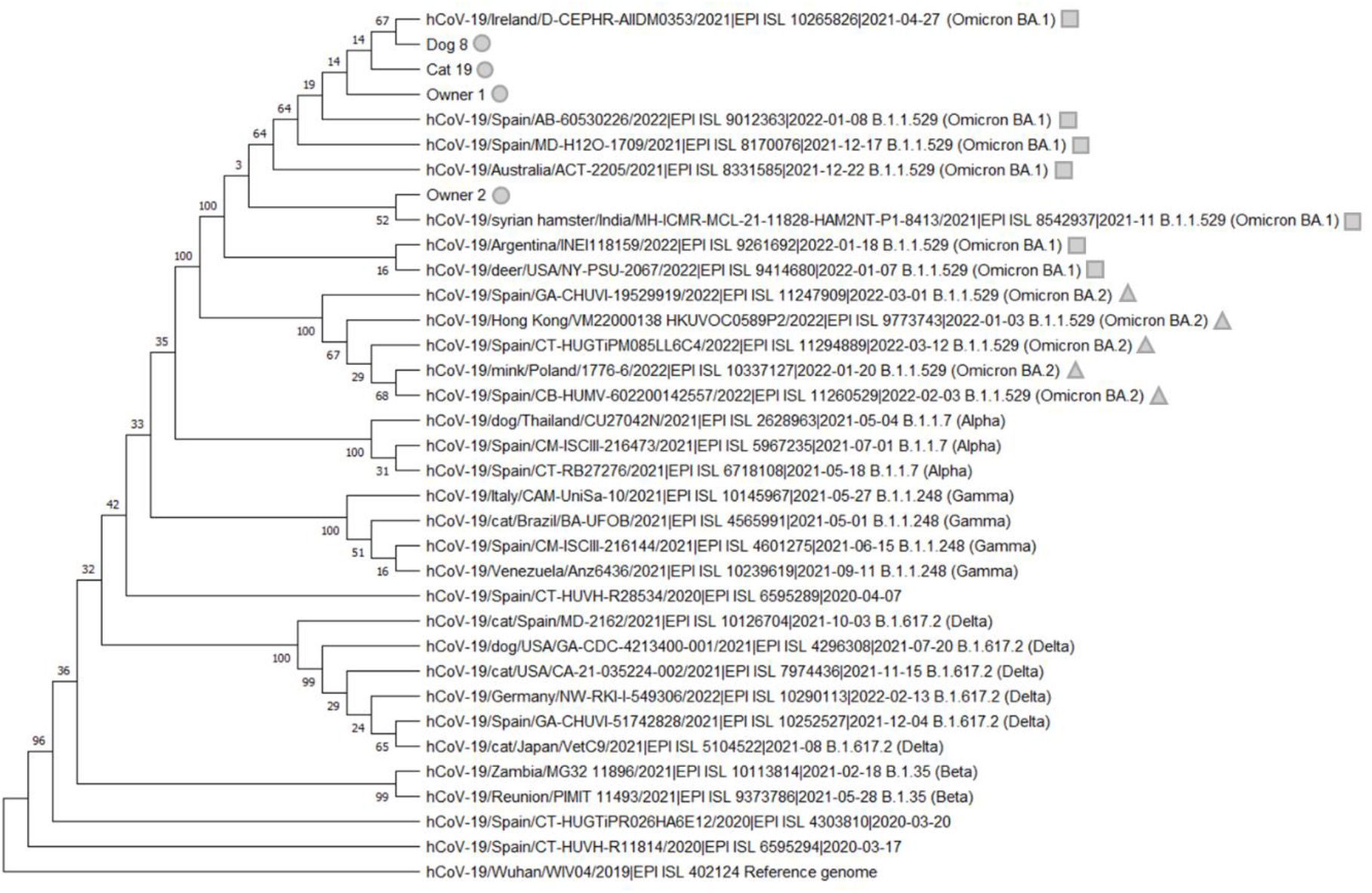
Phylogenetic analysis of SARS-CoV-2 of the whole-genome sequences from Dog_8, Cat_19, Owner_1, and Owner_2 (grey circle), which were clustered with the SARS-CoV-2 B.1.1.529 (Omicron) and more specifically with lineage BA.1 (grey square). The lineage BA.2 is indicated with a grey triangle.

Analysis in the CoVsurver mutations app (GISAID) showed that the sequences presented several mutations having as a reference the hCoV-19/Wuhan/WIV04/2019 sequence. The mutations were 37 in the case of Dog_8 and Cat_19 (Table 2, Table 3). No variabilities were observed at the nucleotide/amino acid level between the sequences from Dog_8 and its owner (Owner_1).

**Table 2.** List of mutations displayed in the different regions of the genome of SARS-CoV-2 in the sequence obtained in this study of Dog_8. NSP: Non-structural protein; E: envelope protein; M: Membrane protein; N: Nucleocapside protein.

| Location in the genome | Mutations displayed |
| --- | --- |
| NSP3 (ORF1a) | K38R, P985S, V1069I, S1265del, L1266I, A1892T |
| NSP4 (ORF1a) | T492I |
| NSP5 | P132H |
| NSP6 (ORF1a) | L105del, S106del, G107del, I189V |
| NSP12 (ORF 1b) | P323L |
| NSP14 (ORF 1b) | I42V |
| Spike | A67V, H69del, V70del, T95I, G142D, V143del, Y144del, Y145del, N211del, L212I, ins214EPE, G339D, S371L, S373P, S375F, K417N, N440K, G446S, S477N, T478K, E484A, Q493R, G496S, Q498R, N501Y, Y505H, T547K, D614G, H655Y, N679K, P681H, N764K, D796Y, N856K, Q954H, N969K, L981F |
| E | T9I |
| M | D3G, Q19E, A63T |
| N | P13L, E31del, R32del, S33del, R203K, G204R |

**Table 3.** List of mutations displayed in the different regions of the genome of SARS-CoV-2 in the sequence obtained in this study of Cat_19. NSP: Non-structural protein; NS3: Non-structural protein 3; E: envelope protein; M: Membrane protein; NS7a: Accessory protein 7a; N: Nucleocapside protein.

| Location in the genome | Mutations displayed |
| --- | --- |
| NSP3 (ORF1a) | K38R, P985S, V1069I, S1265del, L1266I, A1892T |
| NSP4 (ORF1a) | T492I |
| NSP5 | P132H |
| NSP6 (ORF1a) | L105del, S106del, G107del, I189V |
| NSP12 (ORF 1b) | P323L |
| NSP14 (ORF 1b) | I42V |
| Spike | A67V, H69del, V70del, T95I, G142D, V143del, Y144del, Y145del, N211del, L212I, ins214EPE, G339D, S371L, S373P, S375F, K417N, N440K, G446S, S477N, T478K, E484A, Q493R, G496S, Q498R, N501Y, Y505H, T547K, D614G, H655Y, N679K, P681H, N764K, D796Y, N856K, Q954H, N969K, L981F |
| NS3 | T14del, L15del |
| E | T9I |
| M | D3G, Q19E, A63T |
| NS7a | ins45EstopLN |
| N | P13L, E31del, R32del, S33del, R203K, G204R |

## Discussion

The SARS-CoV-2 B.1.529 (Omicron) variant, the last VOC detected, is nowadays highly extended around the world. Concretely in Spain, epidemiological data from the Omicron-associated wave has evidenced that the transmission rate of this variant is quite superior to other variants such as Beta or Delta. This fact has promoted the rapid spread of this variant, being dominant since November 2021 (He, Hong et al. 2021). One concern about this new variant is its potential transmission to other species, in which it could evolve and acquire new mutations that may be involved in higher virulence, among other fears. For this reason, it is necessary to evaluate its capability to infect susceptible species. In this sense, pets such as cats and dogs should be a major focus due to their close contact with humans.

In this study, we evidenced the detection of the Omicron SARS-CoV-2 variant in companion animals, demonstrating that pets are susceptible to the infection with this strain. However, according to the outcomes obtained by this work, there was a relatively low number of positive animals to RT-qPCR taken into account the characteristics of the study, which involved an active sampling. In all the cases, owners assured high contact with their pets. In addition, the sampling was done at the best time for the detection of the disease (Shi, Wen et al. 2020) and only 10.13% of animals became infected, as well as no clinical signs were observed in any animal. These results contrast with previous reports in which the susceptibility of cats and dogs to other SARS-CoV-2 variants such as Alpha and Delta seems to be higher (Barroso-Arévalo, Sánchez-Morales et al. 2022; Barroso-Arevalo, Rivera et al. 2021; Fernandez-Bastit, Rodon et al. 2021, Hamer, Ghai et al. 2021). Furthermore, in the case of animal naturally infected with these other variants, clinical signs were described (Segales, Puig et al. 2020, Ferasin, Fritz et al. 2021, Fernandez-Bastit, Rodon et al. 2021, Hamer, Ghai et al. 2021, Barroso-Arévalo, Sánchez-Morales et al. 2022) and higher viral loads were detected (Barroso-Arevalo, Rivera et al. 2021, Barroso-Arévalo, Sánchez-Morales et al. 2022). Another remarkable difference observed in animals infected with the Omicron variant is that viral isolation was not possible from any sample, due to the low viral load of all positive specimens. The fact that viral isolation was not possible could be due, in part, to the lower fusogenicity of this variant (Suzuki, Yamasoba et al. 2022) which may difficult the virus entry into the cell. By contrast, viral isolation from cat and dogs samples has been possible in the case of the original virus isolate (Hamer, Pauvolid-Correa et al. 2020, Barroso-Arevalo, Barneto et al. 2021) and other variants (Barroso-Arevalo, Rivera et al. 2021). Neither was it possible to detect the presence of neutralizing antibodies in the RT-qPCR-positive animals in contrast to other SARS-CoV-2 seroprevalence studies in animals. This may be derived from the fact that virus replication may have been limited to the local level. In consequence, it is possible viral dissemination did not occur in the infected animals and what we have detected were remnants of viral RNA (Días et al., 2021). This could be explained by the fact that PCR-positive animals were only detected on one day of the 4 to 5 consecutive days of sampling.

All these results may be explained by the fact that a higher affinity with the human cellular receptor has been reported in the case of the Omicron variant compared to other variants (He, Hong et al. 2021, Zou, Xia et al. 2021, Suzuki, Yamasoba et al. 2022). This could have led to the displacement of the binding between the animal cell and the virus, maybe due to specific variations in the ACE2 animal’s receptor with respect to the human ACE2. This may be the reason for the variation of susceptibility in animals to this new variant compared to the previous ones. Further experimental research should be conducted to corroborate this hypothesis since non-experimental data on cats and dogs infected with the Omicron variant are available so far.

However, these results contrast with those of an experimental study carried out in mink (Virtanen, Aaltonen et al. 2022), in which high pathogenicity of the Omicron variant was observed, both at the level of symptoms and post mortem anatomopathological lesions. This higher susceptibility may be affected by the fact that mink-derived SARS-CoV-2 strains encode substitutions in areas of the genome crucial for ACE2 receptor binding that may enhance the binding of the spike protein to this receptor (Welkers, Han et al. 2021). It is, therefore, necessary to carry out studies on the pathogenicity of this variant in different animal species, as well as active surveillance to be able to early detection of new emerging variants.

Although so far there have been no publications on the presence of the Omicron variant in pets, it has been detected in wildlife, specifically in white-tailed deer (*Odocoileus virginianus*), which have been shown to be highly susceptible to the SARS-CoV-2 virus. Fortunately, despite their higher susceptibility, the risk of high contact with an infected human in this species is quite low, contrary to what is happening in the case of pets. These aspects highlight the importance of the investigation of these new variants both in urban and wild fauna (Vandegrift, Yon et al. 2022).

From what we have observed in this study, it appears that the Omicron variant is less virulent to pets than the previous variants as well as the original isolate. Although 10.13% of the animals analyzed in this field study tested positive for RT-qPCR, low viral loads were detected and none of the infected animals showed any symptomatology. This, together with our results previously obtained on VOCs in animals (Barroso-Arevalo, Rivera et al. 2021; Barroso-Arévalo, Sánchez-Morales et al. 2022) has demonstrated the great variability of pathogenicity and response of each animal species and the efficiency of our active surveillance system. This highlights the importance of conducting active surveillance in both pets living with COVID19 infected people and wildlife, in addition to genomic research to early detect infections with other variants or mutations associated with animal hosts. This also underlines the relevance of establishing a network of clinics and owners to be able to carry out active surveillance sampling.

## Acknowledgments and funding

This research was funded by the Institute of Health Carlos III (ISCIII), project “Estudio del potencial impacto del COVID19 en mascotas y linces” (reference: COV20/01385) and the Community of Madrid project “REACT ANTICIPA-UCM” (reference PR38/21).

The authors would like to thank Belén Rivera, Rocío Sánchez and Deborah López for their excellent technical support, as well as all the members of the COVID-VISAVET team. The authors are also grateful to all the Veterinary Clinics and owners who participated in this study, especially Begoña Rodero for her major support.

## Author contributions

SBA and JMSV designed the study. SBA and LSM performed the sampling and veterinary inspection. SBA and LSM performed laboratory analysis. LD and JMSV acquired the funds. SBA and LSM wrote the initial manuscript. LD, MPS, and JMSV reviewed the manuscript. All the authors have read and approved the final version of the manuscript.

## References

Abdelnabi, R., Foo, C. S., Zhang, X., Lemmens, V., Maes, P., Slechten, B., Raymenants, J., André, E., Weynand, B., Dallmeier, K., & Neyts, J. 2022. The omicron (B.1.1.529) SARS-CoV-2 variant of concern does not readily infect Syrian hamsters. Antiviral research, 198, 105253.

Araf, Y., Akter, F., Tang, Y. D., Fatemi, R., Parvez, M., Zheng, C., & Hossain, M. G. 2022. Omicron variant of SARS-CoV-2: Genomics, transmissibility, and responses to current COVID-19 vaccines. Journal of medical virology, 94(5), 1825–1832.

Barroso-Arévalo, S., Barneto, A., Ramos, Á. M., Rivera, B., Sánchez, R., Sánchez-Morales, L., Pérez-Sancho, M., Buendía, A., Ferreras, E., Ortiz-Menéndez, J. C., Moreno, I., Serres, C., Vela, C., Risalde, M. Á., Domínguez, L., & Sánchez-Vizcaíno, J. M. 2021. Large-scale study on virological and serological prevalence of SARS-CoV-2 in cats and dogs in Spain. Transboundary and emerging diseases, 1–16.

Barroso-Arévalo, S., Rivera, B., Domínguez, L., & Sánchez-Vizcaíno, J. M. 2021. First Detection of SARS-CoV-2 B.1.1.7 Variant of Concern in an Asymptomatic Dog in Spain. Viruses, 13(7), 13791.

Barroso-Arévalo, S., L. Sánchez-Morales, M. Pérez-Sancho, L. Domínguez and J. M. Sánchez-Vizcaíno. 2022. First Detection of SARS-CoV-2 B.1.617.2 (Delta) Variant of Concern in a Symptomatic Cat in Spain. Frontiers in Veterinary Science 9, 841430.

Centers for Disease Control and Prevention (US). Science Brief: Omicron (B.1.1.529) Variant. 2021. In National Center for Immunization and Respiratory Diseases (NCIRD), Division of Viral Diseases, CDC COVID-19 Science Briefs.

Chen, J., & Wei, G. W. 2022. Omicron BA.2 (B.1.1.529.2): high potential to becoming the next dominating variant. Research square, rs.3.rs-1362445.

Corman, V. M., Landt, O., Kaiser, M., Molenkamp, R., Meijer, A., Chu, D. K., Bleicker, T., Brünink, S., Schneider, J., Schmidt, M. L., Mulders, D. G., Haagmans, B. L., van der Veer, B., van den Brink, S., Wijsman, L., Goderski, G., Romette, J. L., Ellis, J., Zambon, M., Peiris, M., … Drosten, C. 2020. Detection of 2019 novel coronavirus (2019-nCoV) by real-time RT-PCR. Euro surveillance: bulletin Europeen sur les maladies transmissibles = European communicable disease bulletin, 25(3), 2000045.

Desingu, P. A., & Nagarajan, K. 2022. Omicron BA.2 lineage spreads in clusters and is concentrated in Denmark. Journal of medical virology, 1–5.

Halfmann, P. J., Iida, S., Iwatsuki-Horimoto, K., Maemura, T., Kiso, M., Scheaffer, S. M., Darling, T. L., Joshi, A., Loeber, S., Singh, G., Foster, S. L., Ying, B., Case, J. B., Chong, Z., Whitener, B., Moliva, J., Floyd, K., Ujie, M., Nakajima, N., Ito, M., … Kawaoka, Y. 2022. SARS-CoV-2 Omicron virus causes attenuated disease in mice and hamsters. Nature, 603(7902), 687–692.

Dias, H. G., Resck, M. E. B., Caldas, G. C., Resck, A. F., Da Silva, N. V., Dos Santos, A. M. V., … & Dos Santos, F. B. 2021. Neutralizing antibodies for SARS-CoV-2 in stray animals from Rio de Janeiro, Brazil. PLoS One, 16(3): e0248578.

Doerksen, T., Lu, A., Noll, L., Almes, K., Bai, J., Upchurch, D., & Palinski, R. 2021. Near-Complete Genome of SARS-CoV-2 Delta (AY.3) Variant Identified in a Dog in Kansas, USA. Viruses, 13(10), 2104.

Ferasin L., Fritz M., Ferasin, H., Becquart, P., Corbet, S., Gouilh A, M., Legros V., Leroy, E.M. 2021. Infection with SARS-CoV-2 variant B.1.1.7 detected in a group of dogs and cats with suspected myocarditis. Veterinary Record, 189(9), e944

Fernández-Bastit, L., Rodon, J., Pradenas, E., Marfil, S., Trinité, B., Parera, M., Roca, N., Pou, A., Cantero, G., Lorca-Oró, C., Carrillo, J., Izquierdo-Useros, N., Clotet, B., Noguera-Julián, M., Blanco, J., Vergara-Alert, J., & Segalés, J. 2021. First Detection of SARS-CoV-2 Delta (B.1.617.2) Variant of Concern in a Dog with Clinical Signs in Spain. Viruses, 13(12), 2526.

Gortázar, C., Barroso-Arévalo, S., Ferreras-Colino, E., Isla, J., de la Fuente, G., Rivera, B., Domínguez, L., de la Fuente, J., & Sánchez-Vizcaíno, J. M. 2021. Natural SARS-CoV-2 Infection in Kept Ferrets, Spain. Emerging infectious diseases, 27(7), 1994–1996.

Hamer, S. A., Ghai, R. R., Zecca, I. B., Auckland, L. D., Roundy, C. M., Davila, E., Busselman, R. E., Tang, W., Pauvolid-Corrêa, A., Killian, M. L., Jenkins-Moore, M., Torchetti, M. K., Robbe Austerman, S., Lim, A., Akpalu, Y., Fischer, R., Barton Behravesh, C., & Hamer, G. L. 2021. SARS-CoV-2 B.1.1.7 variant of concern detected in a pet dog and cat after exposure to a person with COVID-19, USA. Transboundary and emerging diseases, 00:1–3.

Hamer SA, Pauvolid-Corrêa A, Zecca IB, Davila E, Auckland LD, Roundy CM, Tang W, Torchetti MK, Killian ML, Jenkins-Moore M, Mozingo K, Akpalu Y, Ghai RR, Spengler JR, Barton Behravesh C, Fischer RSB, Hamer GL. 2021. SARS-CoV-2 Infections and Viral Isolations among Serially Tested Cats and Dogs in Households with Infected Owners in Texas, USA. Viruses. 13(5):938.

He, X., Hong, W., Pan, X., Lu, G., & Wei, X. 2021. SARS-CoV-2 Omicron variant: Characteristics and prevention. MedComm, 2(4), 838–845.

Hu, B., Guo, H., Zhou, P., & Shi, Z. L. 2021. Characteristics of SARS-CoV-2 and COVID-19. Nature reviews. Microbiology, 19(3), 141–154.

Meo, S. A., Meo, A. S., Al-Jassir, F. F., & Klonoff, D. C. 2021. Omicron SARS-CoV-2 new variant: global prevalence and biological and clinical characteristics. European review for medical and pharmacological sciences, 25(24), 8012–8018.

Mohapatra, R. K., Kandi, V., Verma, S., & Dhama, K. 2022. Challenges of the Omicron (B.1.1.529) Variant and Its Lineages: A Global Perspective. Chembiochem: a European journal of chemical biology, e202200059.

Paden, C. R., Tao, Y., Queen, K., Zhang, J., Li, Y., Uehara, A., & Tong, S. 2020. Rapid, Sensitive, Full-Genome Sequencing of Severe Acute Respiratory Syndrome Coronavirus 2. Emerging infectious diseases, 26(10), 2401–2405.

Raj R. 2020. Analysis of non-structural proteins, NSPs of SARS-CoV-2 as targets for computational drug designing. Biochemistry and biophysics reports, 25, 100847.

Saxena, S. K., Kumar, S., Ansari, S., Paweska, J. T., Maurya, V. K., Tripathi, A. K., & Abdel-Moneim, A. S. 2022. Characterization of the novel SARS-CoV-2 Omicron (B.1.1.529) variant of concern and its global perspective. Journal of medical virology, 94(4), 1738–1744.

Segalés, J., Puig, M., Rodon, J., Avila-Nieto, C., Carrillo, J., Cantero, G., Terrón, M. T., Cruz, S., Parera, M., Noguera-Julián, M., Izquierdo-Useros, N., Guallar, V., Vidal, E., Valencia, A., Blanco, I., Blanco, J., Clotet, B., & Vergara-Alert, J. 2020. Detection of SARS-CoV-2 in a cat owned by a COVID-19-affected patient in Spain. Proceedings of the National Academy of Sciences of the United States of America, 117(40), 24790–24793.

Shi, J., Wen, Z., Zhong, G., Yang, H., Wang, C., Huang, B., Liu, R., He, X., Shuai, L., Sun, Z., Zhao, Y., Liu, P., Liang, L., Cui, P., Wang, J., Zhang, X., Guan, Y., Tan, W., Wu, G., Chen, H., … Bu, Z. 2020. Susceptibility of ferrets, cats, dogs, and other domesticated animals to SARS-coronavirus 2. Science (New York, N.Y.), 368(6494), 1016–1020.

Sibai, M., Wang, H., Yeung, P. S., Sahoo, M. K., Solis, D., Mfuh, K. O., Huang, C., Yamamoto, F., & Pinsky, B. A. (2022). Development and evaluation of an RT-qPCR for the identification of the SARS-CoV-2 Omicron variant. Journal of clinical virology: the official publication of the Pan American Society for Clinical Virology, 148, 105101.

Suzuki, R., Yamasoba, D., Kimura, I., Wang, L., Kishimoto, M., Ito, J., Morioka, Y., Nao, N., Nasser, H., Uriu, K., Kosugi, Y., Tsuda, M., Orba, Y., Sasaki, M., Shimizu, R., Kawabata, R., Yoshimatsu, K., Asakura, H., Nagashima, M., Sadamasu, K., … Sato, K. 2022. Attenuated fusogenicity and pathogenicity of SARS-CoV-2 Omicron variant. Nature, 603(7902), 700–705.

Tamura K. 1992. Estimation of the number of nucleotide substitutions when there are strong transition-transversion and G+C-content biases. Molecular biology and evolution, 9(4), 678–687.

Vandegrift, K. J., Yon, M., Surendran-Nair, M., Gontu, A., Amirthalingam, S., Nissly, R. H., Levine, N., Stuber, T., DeNicola, A. J., Boulanger, J. R., Kotschwar, N., Aucoin, S. G., Simon, R., Toal, K., Olsen, R. J., Davis, J. J., Bold, D., Gaudreault, N. N., Richt, J. A., Musser, J. M., … Kuchipudi, S. V. 2022. Detection of SARS-CoV-2 Omicron variant (B.1.1.529) infection of white-tailed deer. bioRxiv. 2022.02.04.479189.

Virtanen, J., K. Aaltonen, K. Kegler, V. Venkat, T. Niamsap, L. Kareinen, R. Malmgren, O. Kivelä, N. Atanasova, P. Österlund, T. Smura, A. Sukura, T. Strandin, L. Dutra, O. Vapalahti, H. Nordgren, R. Kant and T. Sironen. 2022. Experimental infection of mink with SARS-COV-2 Omicron (BA.1) variant leads to symptomatic disease with lung pathology and transmission. bioRxiv. 2022.02.16.480524

Variantes de SARS-CoV-2 en España: Ómicron. 2021, 2022. Centro de Coordinación de Alertas y Emergencias Sanitarias. Ministerio de Sanidad de España. ViralZone, SIB Swiss Institute of Bioinformatics

Welkers, M., Han, A. X., Reusken, C., & Eggink, D. 2021. Possible host-adaptation of SARS-CoV-2 due to improved ACE2 receptor binding in mink. Virus evolution, 7(1), veaa094.

World Health Organization. WHO Health Emergency Dashboard. https://covid19.who.int

Zou, J., Xia, H., Xie, X., Kurhade, C., Machado, R., Weaver, S. C., Ren, P., & Shi, P. Y. 2022. Neutralization against Omicron SARS-CoV-2 from previous non-Omicron infection. Nature communications, 13(1), 852.

